# Distinct roles of directional and positional experience in *de novo* visuomotor learning

**DOI:** 10.64898/2026.01.23.701220

**Authors:** Tomoya Kawano, Shota Hagio

## Abstract

Humans can flexibly acquire entirely new sensorimotor mappings, a process known as *de novo* motor learning. A central challenge in *de novo* motor learning is that the learner must discover a viable solution from scratch within a highly redundant control space, without predefined task constraints. Understanding what types of sensorimotor information contribute to the formation of accurate motor behavior in such situations is therefore critical for explaining how novel sensorimotor skills are acquired. While previous studies have suggested that novel visuomotor mappings can be formed based on movement direction and target position, it remains unclear how these two types of information contribute to the learning process. To address this question, we trained 25 human participants to learn arbitrary joystick-to-cursor mapping. We then employed a generalization paradigm to selectively restrict learning experience to either movement direction or target position. Three distinct target conditions were designed: one emphasized target position (P), another emphasized movement direction (D), and a third (P&D) encouraged learning of both components separately. As a result, direction experience improved movement initiation, whereas position experience enhanced movement termination. However, in the P&D condition, combining these experiences did not yield additive generalization. Instead, endpoint accuracy was positively correlated with the degree of alignment between direction- and position-based joystick outputs within the control space. These results suggest that accurate formation of a novel sensorimotor map depends on the coordinated use of directional and positional experiences.

**Significant Statement:** How do humans build entirely new sensorimotor relationships from scratch? This study examined how distinct sensorimotor experiences (movement direction and target position) contribute to the acquisition of a novel joystick-to-cursor mapping. By isolating these experiences, we found that direction experience improved movement initiation, while position experience enhanced movement termination. However, combining these experiences did not lead to more accurate movements as a whole. Instead, the accuracy was related to how well directional and positional joystick outputs were aligned in a control space. These findings suggest that *de novo* motor learning requires the coordinated use of directional and positional information.

## Introduction

Humans possess a remarkable ability to learn arbitrary sensorimotor maps, a process known as *de novo* motor learning (Krakauer et al., 2019). This capability underlies many everyday skills, such as playing musical instruments, mastering video games, or using unfamiliar tools. For instance, when starting a new video game, players initially have no idea how their controller inputs (motor commands) will translate into the movements of the on-screen character (sensory outcomes). *De novo* motor learning refers to the process of establishing this novel relationship (i.e., forming a new sensorimotor map). Previous studies have primarily focused on motor adaptation, where pre-existing sensorimotor maps are recalibrated in response to environmental changes (Krakauer et al., 2000; Shadmehr and Mussa-Ivaldi, 1994). However, information regarding how such sensorimotor maps are built from scratch remains limited.

To experimentally induce *de novo* motor learning, researchers have devised a variety of tasks. A commonly used approach is to create an arbitrary mapping between the position of a body part and on-screen cursor position (Haith et al., 2022; Kawano et al., 2024; Mosier et al., 2005). For example, in a bimanual task, vertical movements of one hand control horizontal cursor movements and *vice versa*, forcing participants to acquire a novel visuomotor map through practice (Haith et al., 2022; Yang et al., 2021). While participants initially struggle to direct the cursor toward intended targets, they gradually acquire precise control with training. Although these tasks evoke *de novo* motor learning, how the brain constructs a new sensorimotor map remains poorly understood. Unlike motor adaptation, *de novo* motor learning requires establishing entirely new mappings between actions and sensory consequences. This distinction raises a fundamental question: what aspects of the sensorimotor experience does the brain rely on when constructing a novel sensorimotor mapping from scratch?

It has been proposed that the brain can represent visuomotor map based on two distinct types of information (van der Graaff et al., 2017; Wang and Sainburg, 2005; Wu and Smith, 2013). One is to transform a movement direction into a change in hand posture (vector coding); and the other is to transform a target position into a specific hand posture (position coding). Several studies suggested that *de novo* motor learning relies on vector coding (Mosier et al., 2005; Ranganathan et al., 2013). Supporting this idea, it has been observed that the cursor trajectory tended to became straight even without explicit instruction (Mosier et al., 2005). In addition, the final hand posture at the target has been shown to depend on the movement direction toward the target (Ranganathan et al., 2013). Conversely, other findings suggested the involvement of position coding. Several studies have demonstrated that *de novo* motor learning tasks can be learned without continuous visual feedback of a cursor (Kim et al., 2024; Liu and Scheidt, 2008; Park et al., 2024). This finding implies that the brain can construct visuomotor maps using only information about the target position (i.e., position coding). Together, these findings suggest that the brain can build a novel visuomotor map using information about both movement direction and target position.

In this study, we aimed to elucidate how two types of information—movement direction and target position—contribute to *de novo* motor learning. To this end, we employed a generalization paradigm (Donchin et al., 2003; Malfait et al., 2002; Shadmehr, 2004), which allows us to dissociate the contributions of direction and position experiences by examining how restricted learning experience generalizes to untrained movements. In the experiment, three distinct target conditions were designed to elicit generalization from experience with specific components: one target emphasized target position, another emphasized movement direction, and a third encouraged learning of both elements separately. We hypothesized that if vector and position coding additively contribute to the acquisition of a *de novo* sensorimotor map, learning both components separately would lead to greater generalization than learning either component alone.

## Materials and Methods

### Participants

Twenty-five right-handed individuals without a history of neurological or motor disorders were included in this study (13 males and 12 females; aged 20.8 *±* 1.87 years, mean *±* standard deviation (SD)). All participants provided written informed consent after receiving a detailed description of the purpose, potential benefits, and risks of the experiment.

One female participant was excluded due to failure to complete the whole task within the 1.5-hour experiment time (complete up to the 12th block out of 20). All procedures used in this study were performed in accordance with the Declaration of Helsinki and were approved by a local institutional ethics committee (approval number: 24-H-15).

### Experimental setup

The participants sat in front of two three-axis joysticks (Model: 9SA50RR-22-47; APEM, France) equipped on an adjustable table with a 27-inch LCD monitor (1,920 *×* 1,080-pixel resolution, 165 Hz refresh rate; ASUS, Taiwan) (Figure 1A). Both joysticks were positioned symmetrically with respect to the participant’s midline, placed 30 cm forward from the face, and separated laterally by 22.5 cm. Participants could not see their hands, and their head motion was limited by a chinrest. Participants manipulated the two joysticks with both hands using three fingers (thumb, index, and middle finger). Each joystick measured 3-axis positions (i.e., X: horizontal, Y: vertical, and Z: roll axis), acquiring a total of 6 input signals in 16-bit at a sampling rate of 50 Hz. We performed preprocessing on these inputs in real time using a custom LabVIEW program as follows: Initially, joystick inputs were normalized such that each axis could vary independently within the range of [*−*1, 1], where *±*1 corresponded to *±*14.4^◦^ tilt for the X- and Y-axes, and *±*27^◦^ rotation for the Z-axis. Thereafter, the five-point moving average was applied to reduce the intrinsic noise of the joysticks. The post-processing 6-dimensional vector ***h***, representing the joystick positions, was linearly mapped onto the lateral and vertical cursor position ***p*** using the equation below (Choi et al., 2020; Mosier et al., 2005; Ranganathan et al., 2013) (Figure 1B):

**Figure 1.**
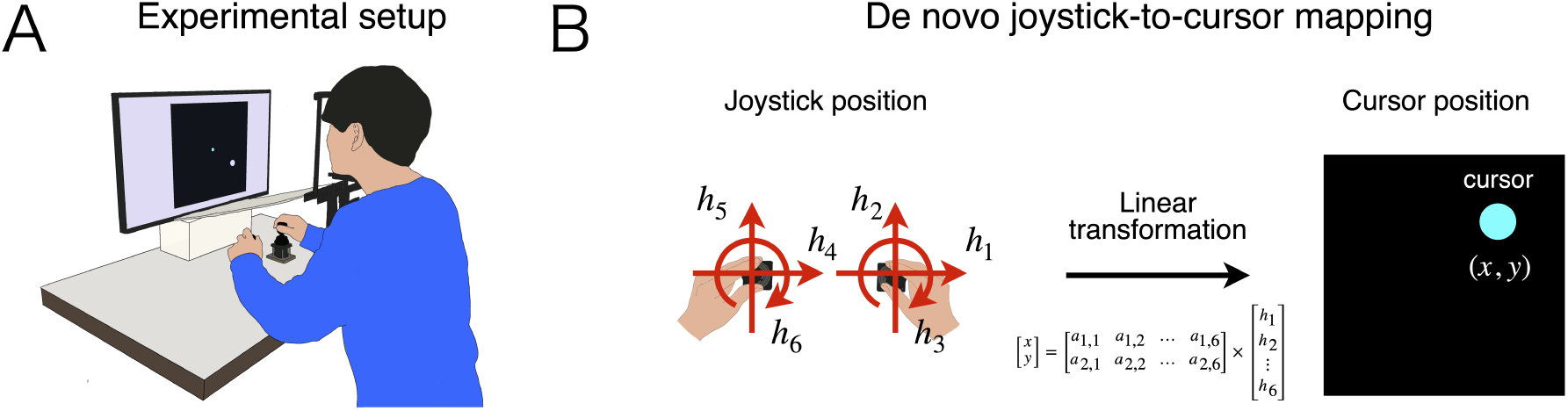
Experimental setup. **A**: Participants manipulated two joysticks to control the on-screen cursor. **B**: Cursor position was determined by the joystick position. Two joysticks measured a total of six input signals. The six-dimensional joystick position vector was linearly translated into the cursor position.

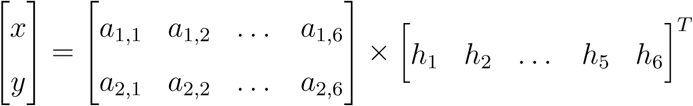

i.e., ***p*** = ***Ah***, where the mapping matrix ***A*** was determined from the calibration task (described below).

### Experimental task

There were two tasks in the experiment: a calibration task and point-to-point reaching task.

#### Calibration task

To determine the mapping matrix ***A*** for each participant, calibration task was conducted. In this task, participants were instructed to move the joysticks freely for 1 minute. To encourage participants to explore various joystick movements, we presented six bar graphs visualizing the real-time variance of joystick positions for each axis (Choi et al., 2020). The acquired calibration data was standardized for each axis to ensure that all axes contributed to the mapping matrix ***A***. Principal component analysis (PCA) was then performed on the standardized calibration data. The first two principal components (PC1 and PC2) were used to form the first and second rows of the mapping matrix ***A***. Therefore, the scores of the PC1 and PC2 were converted to the lateral and vertical cursor position, respectively. We used first two PCs to avoid requiring participants to perform unfamiliar joystick movements in the following reaching task.

#### Point-to-point reaching task

Participants were required to move the on-screen cursor (5 mm radius) to a series of targets (10 mm radius). During the task, targets were presented sequentially, with a sequence of trials referring to a group of consecutive reaches toward these targets (Figure 2A). Details regarding the locations and order of the targets are described below (Experimental conditions). All trials were classified into either a training trial or a test trial. Each trial was defined as one reaching movement toward a single target. In the training trial, participants were given continuous visual feedback of the cursor. Based on this visual information, participants moved the cursor toward the target. When the cursor stayed within a target for 500 ms, it moved to a new location. At the end of every few training trials (referred to as a sequence of trials), the visual feedback of the cursor was removed and participants were instructed to relax the joysticks (i.e., release their hands so that the spring force passively returned each joystick to the center position) to reset the cursor position at the center of the workspace (Figure 2B). To control the amount of time spent across participants, a time limit of 12 seconds was imposed only on the final trial of each sequence. If the elapsed time exceeded 12 seconds, the trial was terminated and participants were instructed to relax the joysticks. Participants were instructed to move the cursor to the target as fast as possible. To motivate them to follow this instruction, we provided a score calculated based on the elapsed time after the start of the trial. The score was calculated as follows.

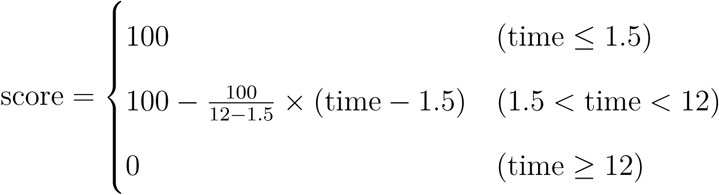

**Figure 2.**
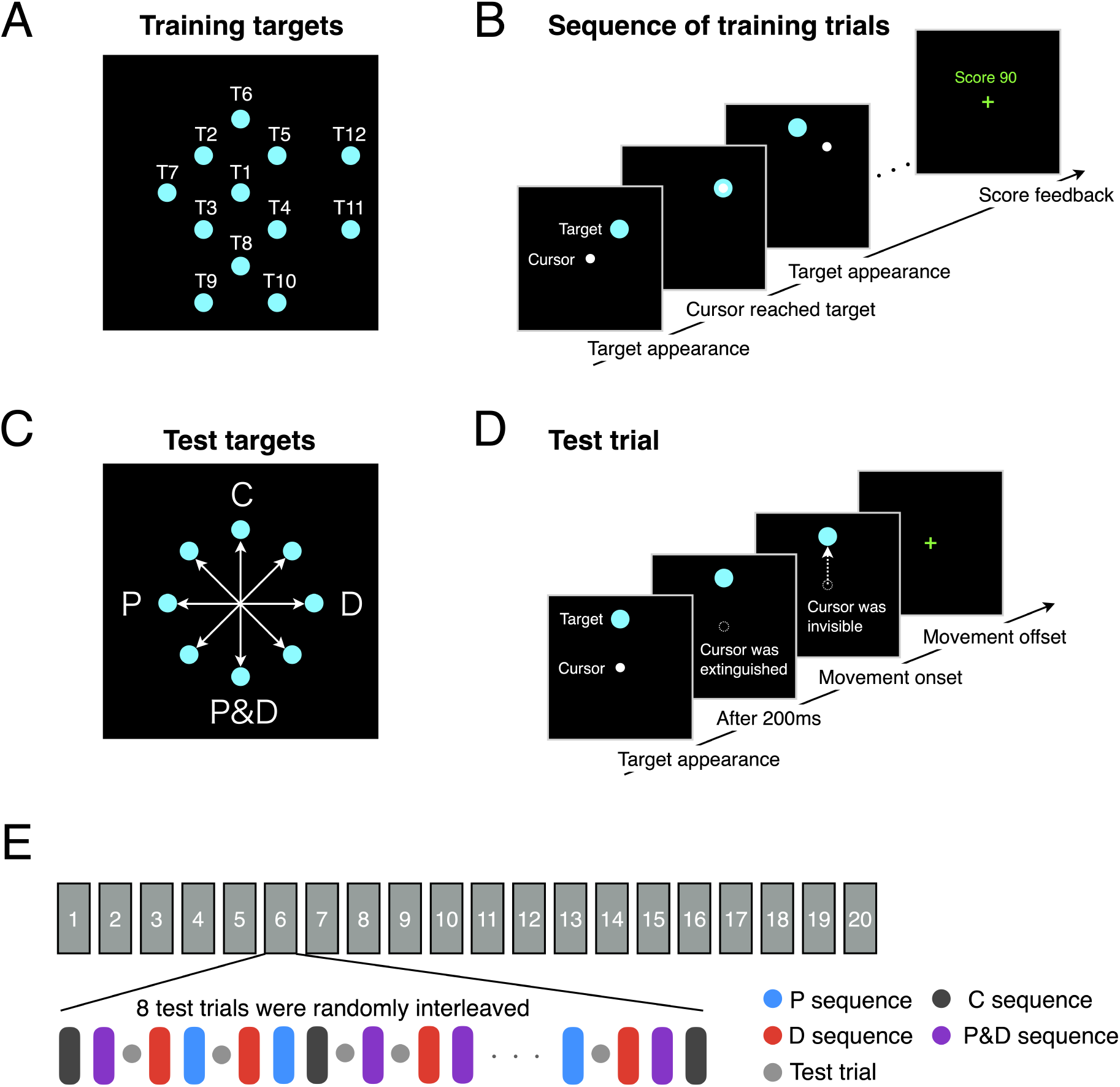
Experimental design. **A**: The target presented in training trials consist of 12 targets, labeled T1–T12. This configuration illustrates one example of the possible arrangements. **B**: The participants performed sequence of training trials to learn visuomotor maps. **C**: The test targets were located in eight directions at intervals of 45^◦^, and the start position was the center of the workspace. **D**: The participants performed test trials to help measure the amount of visuomotor map learning. **E**: The task consists of 20 blocks. In each block there are 20 sequences of training trials (rounded rectangle) and 8 test trials (gray circle). The sequences for each condition were presented in a pseudorandom order every four sequences. In block 1, test trials were presented at the beginning of the block. In the subsequent blocks, test trials were randomly interleaved between sequences. The four colors in the sequences (rounded rectangle) represent the differences in experimental conditions (black: Control, blue: Position, red: Direction, purple: Position and Direction).

In the test trial, the visual feedback of the cursor was removed to evaluate the openloop movement learned during the training trial. The test trials were always initiated from the center of the workspace and directed toward one of the eight targets (Figure 2C). The procedure for the test trial was as follows (Figure 2D):

1. Participants positioned the cursor in the center of the workspace.
2. A target appeared, and 200 ms later the visual feedback of the cursor was removed. The 200 ms is the minimum reaction time for the visuomotor system (Georgopoulos et al., 1981; Soechting and Lacquaniti, 1983; van Sonderen et al., 1988). This duration was set to allow participants to recognize the spatial relationship between the cursor and the target while preventing them from seeing the cursor movement afterwards.
3. Once the participants started moving the joysticks (onset) and then stopped them (offset), the trial ended. The onset was identified when the joystick tangential velocity (i.e., the time derivative of the 6D joystick position, computed as the Euclidean norm of the difference vector divided by the time interval) exceeded a threshold of 0.5 for 10 consecutive time steps. The offset was defined as the time when the joystick tangential velocity fell below a threshold of 0.5 after the onset. After the offset, participants were instructed to relax the joysticks.
4. The visual feedback of the cursor was restored after the joysticks returned to the center position.

To encourage participants to perform open-loop movement, temporal criteria on their behavior was imposed. If the time between the start of the trial and the offset exceeded 2 seconds, participants were provided with a warning message, “stop a little earlier”. The test targets were located in eight directions at intervals of 45^◦^ and a distance of 8 cm (Figure 2C). Among the eight targets, four targets (0^◦^, 90^◦^, 180^◦^, and 270^◦^) were assigned to four different conditions. The assignment of conditions to these four targets was counterbalanced across participants. These targets were then used to implement the different experimental conditions described below.

### Experimental conditions

We designed four experimental conditions to test how learning generalized across untrained movements. To clarify how movement direction and target position contribute to the acquisition of a *de novo* visuomotor map, we employed a generalization paradigm (Donchin et al., 2003; Malfait et al., 2002; Shadmehr, 2004). This approach makes it possible to disentangle the respective roles of direction and position by testing how learning under restricted experience transfers to novel, untrained movements. Within this framework, we defined four conditions that varied in the type of information provided. The Control (C) condition involved direct practice of the test movement, serving as a baseline. The Position (P) condition exposed participants only to the final position of the test movement, while the Direction (D) condition exposed them only to movement direction. The Position and Direction (P&D) condition combined these two components, allowing participants to experience target position and movement direction separately. Figure 2C shows an example configuration of the test targets for the four conditions. The following explanations will be based on this configuration. There are 12 potential targets that could be presented in training trials (T1–T12 in Figure 2A). Participants were trained to move between these targets.

#### Control condition

In this condition, the movement in the test trial was a movement upward from the center of the workspace (Figure 2C). This condition served as a baseline to assess the amount of learning when participants were directly trained on the test movement. To this end, participants were trained to perform the same upward movement as in the test trial. The training was conducted using a sequence of three trials. Initially, the cursor was located at T1. Then, targets were presented in the sequence of either T2 ***→*** T1 ***→*** T6 or T5 ***→*** T1 ***→*** T6 (Figure 2A). The inclusion of the T2 ***→*** T1 (or T5 ***→*** T1) movements was intended to balance the number of movements toward the targets (T2–T5) surrounding the test target. These consecutive three trials are referred to as the C (Control) sequence.

#### Position condition

In this condition, the movement in the test trial was a movement to the left from the center of the workspace (Figure 2C). Participants were allowed to experience only the target position of the movement. To achieve this, they were trained to move to the same position, but from a different direction. The training was conducted using a sequence of two trials. Initially, the cursor was located at T1. Then, targets were presented in the sequence of either T2 ***→*** T7 or T3 ***→*** T7 (Figure 2A). Importantly, T7 was the same as the test target of this condition (Figure 2C). These consecutive two trials are referred to as the P (Position) sequence.

#### Direction condition

In this condition, the movement in the test trial was a movement to the right from the center of the workspace (Figure 2C). Participants were allowed to experience only the direction of the movement. To this end, they were trained to move from the same direction to different positions. The training was conducted using a sequence of two trials. Initially, the cursor was located at T1. Then, targets were presented in the sequence of either T5 ***→*** T12 or T4 ***→*** T11 (Figure 2A). Crucially, T5 ***→*** T12 and T4 ***→*** T11 movements matched the movement direction of the test trial (Figure 2C). This consecutive two trials are referred to as the D (Direction) sequence.

#### Position and Direction condition

In this condition, the movement in the test trial was a movement downward from the center of the workspace (Figure 2C). Participants were allowed to separately experience both the position and direction of the movement. To this end, participants were trained using both the P- and D sequences. Initially, the cursor was located at T1. Then, targets were presented in the sequence of either T3 ***→*** T8, T3 ***→*** T9, T4 ***→*** T8, or T4 ***→*** T10 (Figure 2A). Essentially, T8 was the same as the test target of this condition. Additionally, T3 ***→*** T9 and T4 ***→*** T10 movements matched the movement direction of the test trial (Figure 2C). These consecutive two trials are referred to as the P&D (Position and Direction) sequence. If the effects of direction and position experience additively contribute to the acquisition of a *de novo* sensorimotor map, learning both components separately would yield greater generalization than learning either component alone (P condition and D condition).

### Experimental procedure

The experimental task consisted of 20 blocks (Figure 2E). Each block consisted of 20 training sequences and 8 test trials (Figure 2E). The sequences for each condition were presented in a pseudorandom order every four sequences. In block 1, test trials were presented at the beginning of the block to test movements with minimal training effects. In the subsequent blocks, test trials were randomly interleaved between sequences. The rest period between blocks was 30 seconds, with a longer 120 seconds rest period provided every five blocks.

### Data analysis

All data were analyzed using MATLAB R2025a (MathWorks, Inc., Natick, MA, USA). We recorded cursor position, joystick position, and joystick tangential velocity at 50Hz. The joystick tangential velocity was low-pass filtered (5 Hz) using a fourth-order Butterworth filter.

#### Task performance

The task performance in the training trials was assessed by movement time. The movement time was the time elapsed from movement onset to the end of the trial. The movement onset was defined as the time when joystick tangential velocity exceeded a threshold of 0.5 for 10 consecutive time steps. In the test trial, we calculated final position error calculated as the Euclidean distance between the final cursor position and the target position. In addition to the accuracy of the final cursor position, we evaluated the accuracy of movement initiation by quantifying the initial direction error. This error was quantified as the absolute angle between the direction from the center of the workspace to the cursor position at the time when the joystick tangential velocity reached its first peak and the direction from the center of the workspace to the target position. The peak of the joystick tangential velocity was identified using the MATLAB function findpeaks, with the argument MinPeakProminence set to 5% of the maximum velocity in each trial.

#### Position- and Direction-based visuomotor outputs in joystick control space

In the P&D condition, the effects of position and direction experiences appear additive in the workspace since the direction experience matches the movement direction of the test trial and the position experience matches the target position (Figure 5A). However, in the joystick control space, which includes task-irrelevant dimensions, visuomotor outputs based on direction and position are not necessarily aligned (Figure 5B). We therefore tested how the degree of alignment between position- and direction-based visuomotor outputs influenced generalization in the P&D condition. To this end, we calculated how target positions or movement directions were transformed into joystick positions or joystick movement directions, respectively. The former was defined as the position-based visuomotor output (**VO**_pos_), and the latter as the direction-based visuomotor output (**VO**_dir_). To estimate **VO**_pos_, we averaged the final joystick positions across trials directed to the same target position. Similarly, to estimate **VO**_dir_, we calculated joystick movement vectors as the difference between the final and initial joystick positions in each trial, and then averaged these vectors across trials with the same movement direction in the workspace. For this analysis, we used training trials from the last five blocks. Because the number of trials for averaging was small (12 or 13), a bootstrap procedure with 1,000 resamples was used to obtain stable estimates of these averages. Finally, to quantify the alignment between the two outputs, we calculated *ǁ***VO**dir *−* **VO**pos*ǁ*, where a smaller value indicates greater alignment.

### Outlier criteria

To avoid outlier influenced bias during hypothesis testing, outliers were removed using the quartile method. First, the first quartile (Q1) and the third quartile (Q3), which represent the 25th and 75th percentiles of the data, respectively, were calculated. Thereafter the interquartile range (IQR) was calculated by subtracting Q1 from Q3:

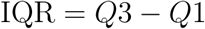

Next, the lower- and upper bound for outliers were calculated:

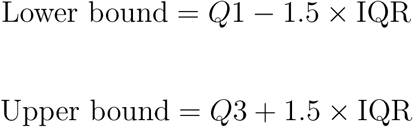

Any data points outside this range were considered outliers.

### Statistics

All statistical analyses were calculated using R version 4.2.2 (Rstudio, Inc., Boston, MA, USA). Data from 24 participants were analyzed. To investigate the effects of condition and trial on the error in last 5 test trials (either final position error or initial direction error), we conducted a linear mixed-effects model analysis using the lme4 package in R (Bates et al., 2015). First, outliers within each condition were excluded (A total of 2.1% and 4.6% of the data was removed for final position error and initial direction error, respectively). After outlier removal, the analyses included 470 observations for final position error and 463 observations for initial direction error. The fixed effects included condition and trial. The random effects included both random intercepts and random slopes for participants. Statistical significance of the fixed effects was determined using Type III *F* -tests. Post hoc pairwise comparisons were conducted using t-tests on estimated marginal means derived from the linear mixed-effects models (using the emmeans package in R), and *p*-values were adjusted for multiple comparisons using the Benjamini-Hochberg procedure. In all statistical analyses, a *p*-value of less than 0.05 was considered statistically significant. Cohen’s d was computed as effect size. An overview of the data structure and statistical tests used in this study is provided in Table 1.

**Table 1:** Statistical table summarizing data structure and statistical tests.

|  | **Data structure** | **Type of test** | **95% CI** |
| --- | --- | --- | --- |
| a | Normal distribution | Type III ANOVA | – |
| b | Normal distribution | Type III ANOVA | – |
| c | Normal distribution | Post hoc pairwise $t$-test | $[-16.7, -1.46]$ |
| d | Normal distribution | Post hoc pairwise $t$-test | $[-1.53, 15.6]$ |
| e | Normal distribution | Type III ANOVA | – |
| f | Normal distribution | Type III ANOVA | – |
| g | Normal distribution | Post hoc pairwise $t$-test | $[-2.02, -0.137]$ |
| h | Normal distribution | Post hoc pairwise $t$-test | $[-2.48, -0.564]$ |
| i | Normal distribution | Post hoc pairwise $t$-test | $[-2.30, -0.560]$ |
| j | Normal distribution | Post hoc pairwise $t$-test | $[-1.08, 1.27]$ |
| k | Normal distribution | Post hoc pairwise $t$-test | $[-1.21, 0.507]$ |
| l | Normal distribution | Pearson correlation | $[0.108, 0.746]$ |

## Results

### Task performance improved in the training trials

Participants learned the *de novo* visuomotor task through repeated training. Figure 3A shows the cursor trajectories of a representative participant in training trials. In the early stages, the cursor movements were erratic under all conditions. In contrast, in the later stages, the cursor moved directly toward the target more efficiently. The movement time decreased as the training progressed, indicating that participants learned to move the cursor to the target more quickly (Figure 3B).

**Figure 3.**
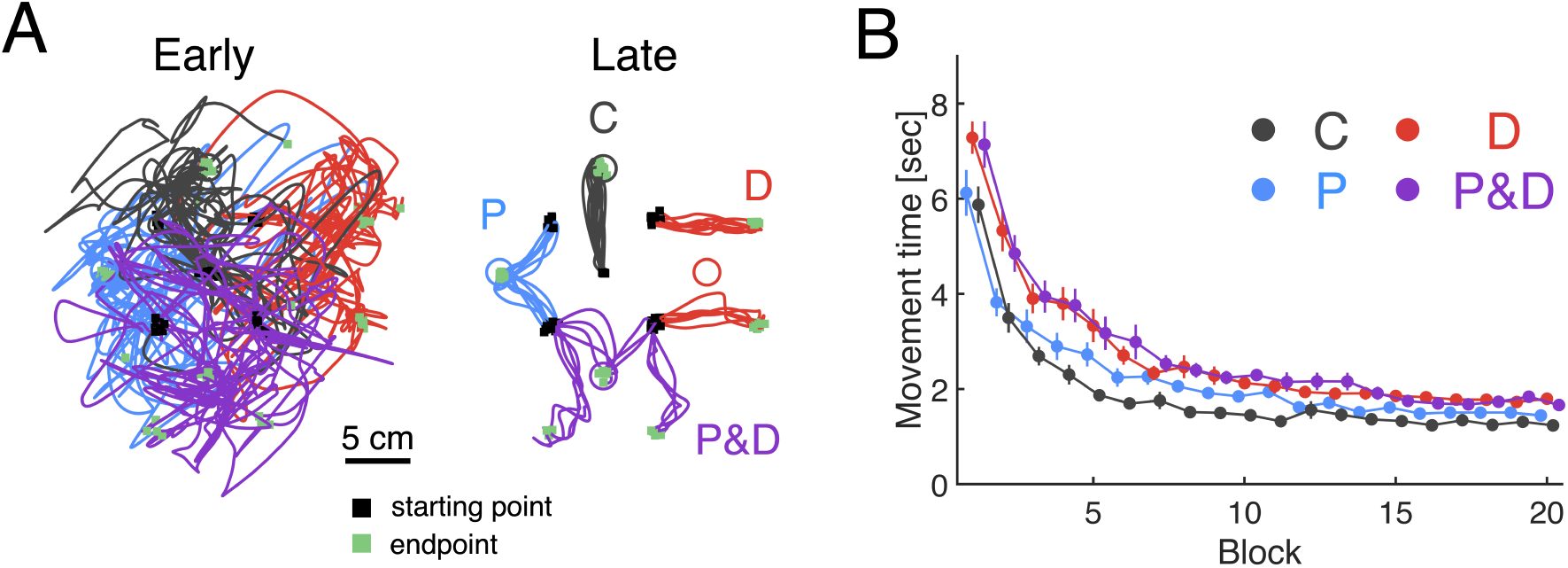
Raw cursor trajectories and Learning curve. **A**: The raw cursor trajectories of a representative participants during training trials in the early stages (first 15 trials for each condition) and the late stages (last 15 trials for each condition). The four colors represent the differences in experimental conditions (black: Control, blue: Position, red: Direction, purple: Position and Direction). For each condition, the final trial of each sequence is displayed. The black square and green square represent the starting point and the endpoint of the cursor trajectory, respectively. **B**: Movement time across blocks. Thick lines indicate the mean across participants. Error bars indicate mean *±* standard error across participants.

### Generalization in the test trials

During the course of training, participants performed test trials on eight targets. Among the eight targets, four targets were assigned to four different conditions. Each condition was different in terms of the experience of movement direction and target position. To assess the ability to reach the target in the test trials, we quantified two metrics, initial direction error and final position error. Through training, both errors in the test trials decreased across trials (Figure 4A and D). Because the test trials were performed with-out online visual feedback, this reduction reflects improvements in feedforward control. Notably, despite the lack of direct training, error reduction was observed in the P, D, and P&D conditions, indicating that generalization had occurred (Figure 4A and D).

**Figure 4.**
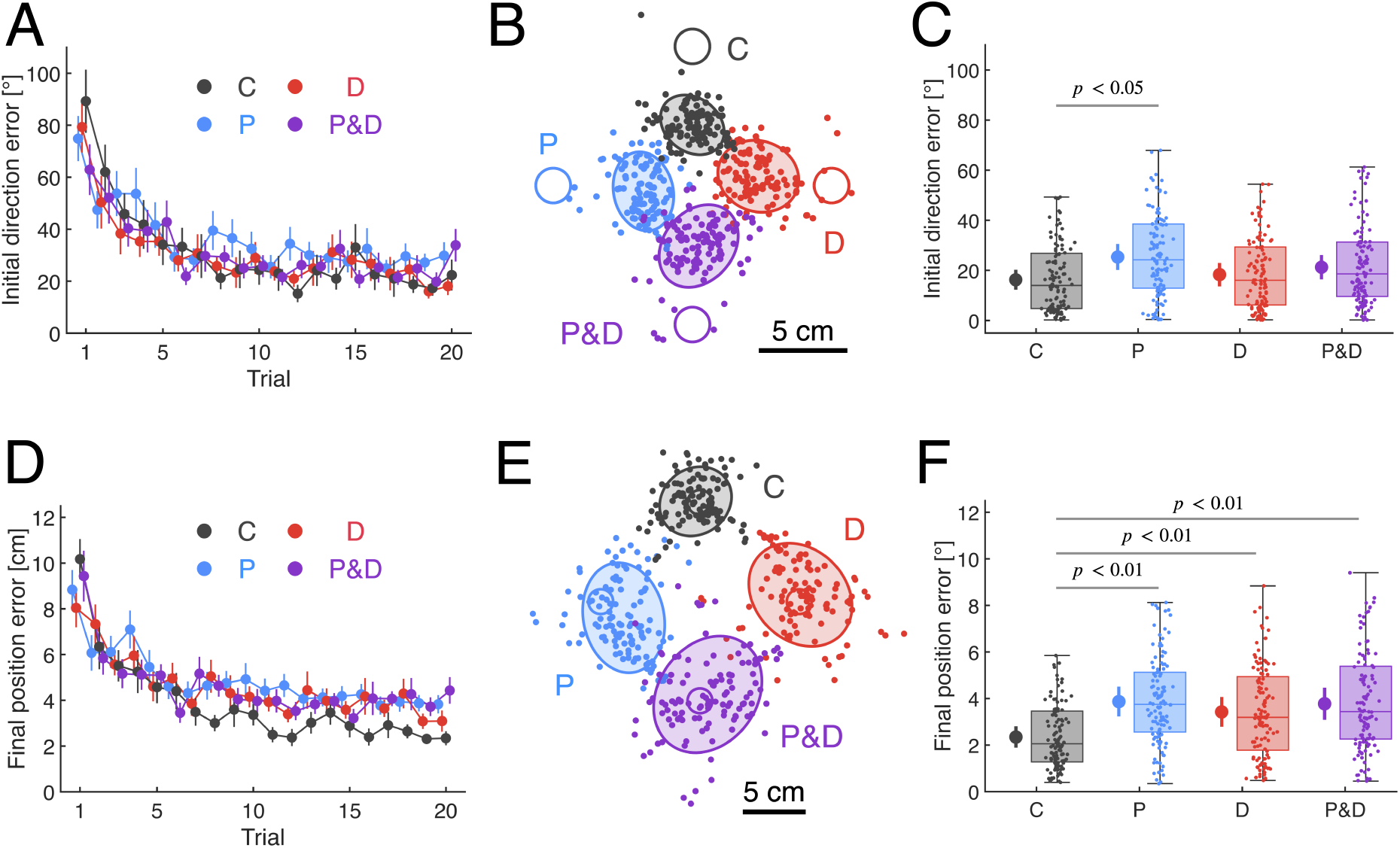
Initial direction error and final position error. **A**: Initial direction error across test trials. Thick lines indicate the mean across participants. Error bars indicate mean *±* standard error across participants. **B**: Distribution of cursor positions at movement initiation. The dots represent the cursor position in each trial pooled across all participants (removing outliers using the Inter quartile range (IQR) method). Ellipses indicate 68% confidence intervals. **C**: The boxplot of initial direction error in the last 5 test trials. The dots represent the error of each trial pooled across all participants. The points next to the boxplots represent the the marginal means for each condition estimated by a linear mixed-effects model. The error bars represent the 95% confidence intervals of the marginal means. **D**: Final position error across test trials. Thick lines indicate the mean across participants. Error bars indicate mean *±* standard error across participants. **E**: Distribution of cursor positions at movement termination. The dots represent the cursor position in each trial pooled across all participants (removing outliers using the IQR method). Ellipses indicate 68% confidence intervals. **F**: The boxplot of final position error in the last five test trials. The dots represent the error of each trial pooled across all participants. The points next to the boxplots represent the the marginal means for each condition estimated by a linear mixed-effects model. The error bars represent the 95% confidence intervals of the marginal means.

### Contribution of direction experience to movement initiation

Although generalization occurred in untrained conditions, the pattern of errors varied depending on the condition. We first examined the initial movement trajectory in the test trials. Figure 4B shows the distribution of the cursor position when the joystick velocity reached its peak. In particular, the distributions differed in shape between the P and D conditions. In the D condition, the distribution was elongated along the target direction, whereas in the P condition, it was elongated perpendicular to the target direction. This suggests that movement initiation in the D condition was aligned with the target direction. These findings were supported by a quantitative comparison of the initial direction error across conditions (Figure 4C). Statistical analysis showed that there was no significant effect of trial, indicating that the learning reached plateau (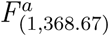 = 0.459, *p* = 0.498). In contrast, there was a significant main effect of condition (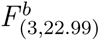 = 4.00, *p* = 0.0199). Post hoc pairwise comparisons revealed that the initial direction error in C condition was significantly lower than in the P condition (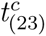 = *−*3.44, *p* = 0.0134, *d* = *−*0.840, 95% CI [*−*16.7, *−*1.46]). This result suggest that the lack of direction experience impaired generalization. Furthermore, there was a weak trend that the initial direction error in D condition was smaller than P condition (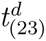 = 2.37, *p* = 0.0794, *d* = 0.651, 95% CI [*−*1.53, 15.6]). Altogether, these results suggest that the experience with movement direction contributed to improved accuracy of movement initiation.

### Contribution of position experience to movement termination

To evaluate the accuracy of movement termination, we analyzed the final cursor position in the test trials. Figure 4E shows the distribution of the final cursor position across conditions. Similar to the movement initiation, the distributions of final positions differed across conditions. In the D condition, although the distribution was centered on the target, frequent overshooting occurred. In contrast, in the P condition, although the center of the distribution was shifted away from the target, overshooting trials were much less frequent. We compared the final position error across conditions. Statistical analysis showed that there was no significant effect of trial, indicating that the learning converged (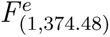 = 0.136, *p* = 0.713). However, a significant main effect of condition (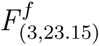 = 12.3, *p* = 5.02 *×* 10^−5^) was observed. Post hoc pairwise comparisons showed that the final position error in D condition was significantly higher than C condition (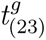 = *−*3.31, *p* = 0.0062, *d* = *−*0.848, 95% CI [*−*2.02, *−*0.137]), suggesting that position experience is necessary for accurately planning where to stop a movement. Furthermore, the final position error in the P condition was also significantly higher than in the C condition (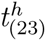 = *−*4.59, *p* = 0.0004, *d* = *−*1.20, 95% CI [*−*2.48, *−*0.564]), indicating that position experience alone is insufficient for accurate movement termination. This result implies that direction experience is a prerequisite for effectively utilizing position experience. Taken together, these findings suggest that position and direction experience interactively contribute to the acquisition of precise movement termination.

### Do position and direction experiences additively contribute to performance?

Given the interactive effects of position and direction experience on movement termination, we then evaluated whether exposing participants to direction and position separately could improve performance. To address this question, we designed a P&D condition in which participants experienced position and direction separately. As a result, statistical analysis revealed that the final position error in the P&D condition was significantly higher than C condition (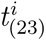 = *−*4.75, *p* = 0.0004, *d* = *−*1.12, 95% CI [*−*2.30, *−*0.560]). In addition, there were no significant differences between P&D vs P (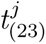 = 0.230, *p* = 0.820, *d* = 0.0734, 95% CI [*−*1.08, 1.27]), P&D vs D (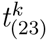 = *−*1.18, *p* = 0.351, *d* = *−*0.275, 95% CI [*−*1.21, 0.507]), suggesting that learning both components separately did not yield greater generalization than learning either component independently. Altogether, these results indicate that experiencing position and direction separately does not necessarily improve the accuracy of movement termination.

### Additive effects depend on visuomotor outputs alignment in control space

In the P&D condition, additive effects were not generally observed, but some participants exhibited these additive effects, suggesting that additive effects may emerge under certain situations (Figure 4F). In the workspace, position and direction experiences appear additive, since the direction experience matches the test movement direction and the position experience matches the test target position (Figure 5A). However, in the joystick control space, there are dimensions that do not affect the resulting cursor movement (task-irrelevant dimensions). Consequently, direction- and position-based motor visuomotor outputs do not necessarily align (Figure 5B). This led to the hypothesis that the additive benefit of P&D experience depends on the alignment of these outputs in the control space. To test this, we quantified the alignment between the two outputs. As a result, participants with higher alignment exhibited smaller final position errors in the P&D condition (*r^l^* = 0.490, *p* = 0.0150, 95% CI [0.108, 0.746]; Figure 5C). These results suggest that the alignment of direction- and position-based visuomotor outputs in the control space may underlie more precise movement control at the endpoint.

**Figure 5.**
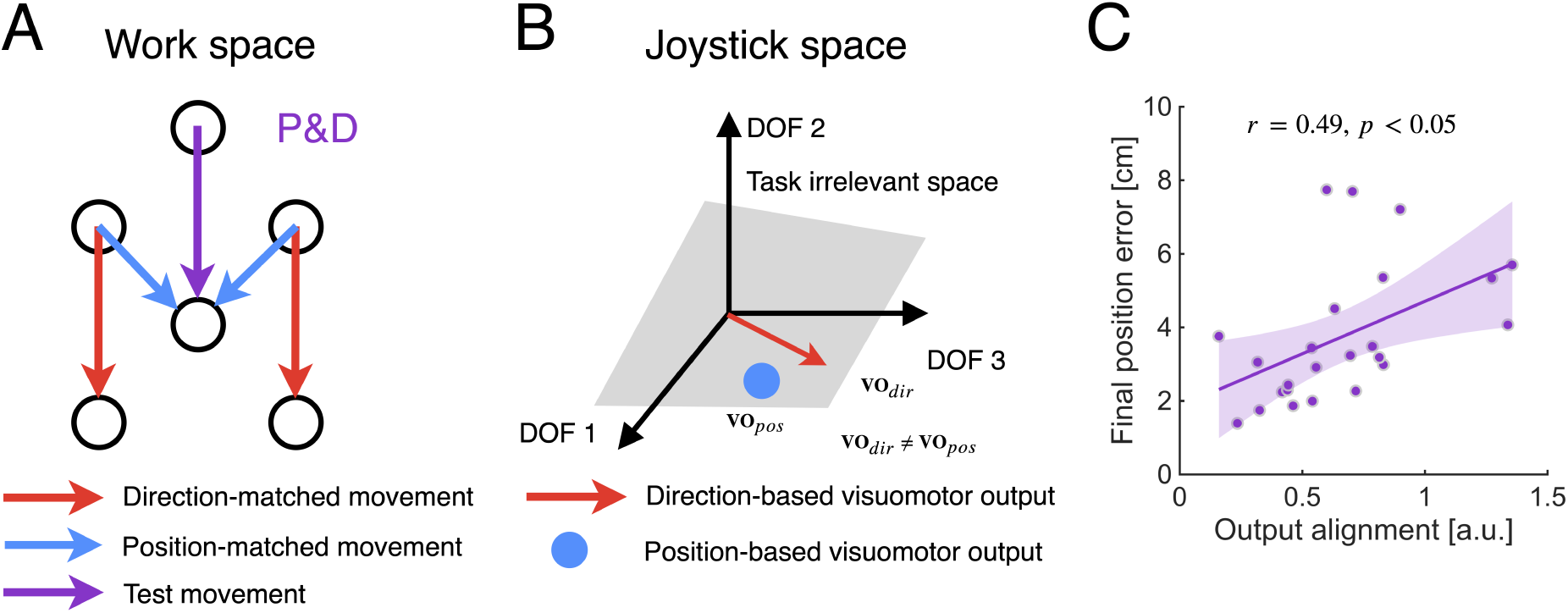
Alignment of two visuomotor outputs in the joystick control space. **A**: The purple arrow indicates the test movement in the Position and Direction (P&D) condition. During training, participants practiced the red movement (direction-matched) and the blue movement (position-matched). The red and purple arrows share the same movement direction, while the blue and purple arrows share the same end position. Thus, in the workspace, it appears as if the direction experience (red) and position experience (blue) could be additively combined to produce the test movement (purple). **B**: Task-irrelevant dimensions exist in the joystick control space (gray plane). As a result, the direction-based motor plan **VO**_dir_ and the position-based visuomotor outputs **VO**_pos_ do not necessarily coincide (**VO**_dir_ */*= **VO**_pos_). The figure illustrates that these two visuomotor outputs are misaligned in the control space, emphasizing that a simple vector addition of direction and position experiences does not necessarily reproduce the test movement. **C**: The scatter plot shows the relationship between output alignment (x-axis) and final position error in the P&D condition (y-axis). Each dot represents an individual participant, the solid line indicates the regression line, and the shaded area denotes its 95% confidence interval.

## Discussion

This study aimed to clarify how two types of information, movement direction and target position, contribute to *de novo* motor learning. To this end, we adopted a generalization paradigm to dissociate the respective contributions of direction and position experiences (Donchin et al., 2003; Malfait et al., 2002; Shadmehr, 2004). The results suggested that direction experience enhanced the accuracy of movement initiation, whereas position experience supported accurate movement termination (Figure 4). However, experiencing position and direction separately did not yield additive effects (Figure 4F). Instead, the study found that participants whose position- and direction-based visuomotor outputs were more closely aligned in the control space exhibited smaller final position errors (Figure 5C).

### Contribution of direction experience to movement initiation

Our results demonstrate that directional experience selectively enhances the accuracy of movement initiation (Figure 4C), supporting the long-standing view that motor learning primarily relies on direction-based representations. This view is supported by classical findings in motor adaptation showing that adaptation to visuomotor rotations involves the remapping of movement vectors rather than final limb positions (Wang and Sainburg, 2005). Evidence from *de novo* motor learning studies further supports this direction-based representation. When participants acquired novel hand-to-cursor mappings, their trajectories became progressively straighter with practice, indicating the establishment of a direction-based transport map linking changes in hand posture to cursor motion (Mosier et al., 2005). Moreover, Ranganathan et al. (2013) demonstrated that the motor system exploits redundancy not by converging to a single final posture, but by adopting distinct hand postures for a single target location, depending on the starting hand configuration. This dependence on the initial state reinforces the dominance of a transport map, i.e., associating cursor displacement with changes in hand posture (Ranganathan et al., 2013). Consistent with these studies, our results showed that isolated experience of movement direction improved the accuracy of movement initiation to a level comparable to that achieved through direct practice (Figure 4C). Altogether, these findings suggest that directional experience contributes to the refinement of direction-based control mechanisms that support accurate movement initiation.

### Contribution of position experience to movement termination

Although initial direction error in the Direction (D) condition was comparable to the Control (C) condition, the final position error was significantly larger in the D condition, indicating impaired accuracy in movement termination (Figure 4). This dissociation can be explained by the existence of two complementary control processes: a transient movement controller that generates vectorial commands for reaching, and a sustained hold controller that stabilizes the limb at the desired endpoint (Albert et al., 2020; Shadmehr, 2017). Evidence for this distinction comes from multiple lines of research. In adaptation studies, learning acquired in visuomotor rotation tasks generalizes poorly between trajectory-reversal (slicing) and positioning (reaching) tasks, suggesting that initial trajectory and final position rely on separate neural mechanisms (Ghez et al., 2007; Scheidt and Ghez, 2007). Neurophysiological data further indicates that neurons in the primary motor cortex exhibit separate load-related tuning during posture and movement, reinforcing the functional separation between control phases (Kurtzer et al., 2005). Accordingly, the impaired termination observed in the D condition likely reflects insufficient formation of the hold controller which depends on positional experience. In addition, the Position (P) condition, in which such positional experience was available, showed improved stabilization but still exhibited larger endpoint errors than the C condition. This indicates that positional experience alone is insufficient, and that the hold controller depends on a well-learned movement controller. Taken together, these findings suggest that positional experience contributes to the learning of the hold controller, which governs accurate movement termination through its interactive contribution with the movement controller.

### Additivity of direction and position experience

One key question was whether separating direction and position experience would additively contribute to generalization when combined (P&D condition). Although additive benefits were not observed at the group level, participants with higher alignment between direction- and position-based visuomotor outputs in the joystick control space exhibited smaller final position errors (Figure 5C). These findings can be interpreted within the framework of the movement and hold controllers (Albert et al., 2020; Shadmehr, 2017). Given the redundancy inherent in the control space, the state reached by the movement controller does not necessarily coincide with the stable state specified by the hold controller. When the misalignment between these states is large, the hold controller may fail to effectively stabilize the joysticks at movement termination. This misalignment may therefore account for the individual variability in final position error observed in the P&D condition. Although previous studies established the existence of distinct movement and hold controllers, how these controllers are learned to support accurate reaching remained unclear (Albert et al., 2020; Shadmehr, 2017). The present findings suggest that accurate reaching depends not only on acquiring the two controllers, but also on coordinating them by aligning the states specified by each controller. An open question remains whether this coordination occurs passively or whether it can be actively learned, and why such individual differences occur.

### Distinction between *de novo* motor learning and motor adaptation

*De novo* motor learning fundamentally differs from motor adaptation in that it necessitates establishing entirely new mappings between motor commands and sensory outcomes, rather than merely recalibrating pre-existing ones. Although prior motor adaptation studies have revealed the existence of separate movement and hold controllers (Ghez et al., 2007; Scheidt and Ghez, 2007), the specific sensorimotor experiences required for their initial construction remain largely unexplored within that traditional paradigm. By utilizing *de novo* motor learning, we were able to precisely control the available experiences, revealing that different types of experience support the acquisition of these two distinct controllers. A second key distinction lies in the treatment of redundancy. Motor adaptation typically involves recalibration within a constrained control space (Berger et al., 2022, 2013; Sadtler et al., 2014), whereas *de novo* motor learning requires the exploration of solutions in an unconstrained high-dimensional space (Choi et al., 2020; Dal’Bello and Izawa, 2021; Mosier et al., 2005). This fundamental difference introduces a distinct computational challenge. When movement and hold controllers are constructed independently in a high-dimensional space, the motor commands specified by each controller are not guaranteed to be mutually consistent. As a result, the state reached by the movement controller may not coincide with the state stabilized by the hold controller, leading to ineffective movement termination. Consequently, in *de novo* motor learning, achieving coordination between these newly acquired controllers emerges as a critical computational requirement for accurate reaching movements.

### Neural representations underlying direction- and position-based learning

To independently achieve direction- and position-based learning, the brain must represent the target in two distinct ways: the cursor-centered displacement vector that specifies movement direction, and the absolute target position that anchors behavior in external space. These two forms of information correspond naturally to the distinction between allocentric and egocentric spatial representations (Chen and Crawford, 2020; Milner and Goodale, 2008; Schenk, 2006). Humans have been shown to reach and point most accurately when both egocentric and allocentric cues are available, suggesting that the visuomotor system benefits from maintaining multiple spatial representations in parallel (Hay and Redon, 2006; Krigolson and Heath, 2004). Neurophysiological evidence further indicates that these two frames of reference are supported by partially distinct cortical systems. Egocentric representations encode the location of the target relative to the viewer and are primarily supported by regions within the dorsal visuomotor stream, including parietal and frontal areas that maintain target position with respect to the self (Chen et al., 2014). In contrast, allocentric representations encode spatial relations relative to external landmarks and rely on the ventral visual stream, particularly occipital-temporal regions involved in landmark-based spatial processing (Chen et al., 2014). Because these two reference frames are implemented by dissociable neural systems, egocentric and allocentric information could be maintained and updated in parallel (Byrne and Crawford, 2010). From this perspective, the two forms of learning observed in our study map naturally onto these distinct spatial codes: position-based learning reflects reliance on egocentric representations of the target location, whereas direction-based learning reflects reliance on allocentric representations of the spatial relation between the cursor and the target. The coexistence of these learning processes is therefore consistent with the view that the brain maintains multiple spatial representations to support flexible visuomotor behavior.

## Conclusion

In conclusion, our results show that *de novo* motor learning relies on both movement direction and target position experiences. Direction experience primarily guides movement initiation, whereas position experience ensures accurate movement termination. Furthermore, when direction and position experiences were provided separately, participants whose direction- and position-based visuomotor outputs were more closely aligned in the control space exhibited higher endpoint accuracy. These findings suggest that accurate formation of a novel sensorimotor maps requires the coordinated use of directional and positional experiences.

